# Predicting drug sensitivity of cancer cells based on DNA methylation levels

**DOI:** 10.1101/2020.08.25.266049

**Authors:** Sofia P. Miranda, Fernanda A. Baião, Paula M. Maçaira, Julia L. Fleck, Stephen R. Piccolo

## Abstract

Cancer cell lines, which are cell cultures developed from tumor samples, represent one of the least expensive and most studied preclinical models for drug development. Accurately predicting drug response for a given cell line based on molecular features may help to optimize drug-development pipelines and explain mechanisms behind treatment responses. In this study, we focus on DNA methylation profiles as one type of molecular feature that is known to drive tumorigenesis and modulate treatment responses. Using genome-wide, DNA methylation profiles from 987 cell lines from the Genomics of Drug Sensitivity in Cancer database, we applied machine-learning algorithms to evaluate the potential to predict cytotoxic responses for eight anti-cancer drugs. We compared the performance of five classification algorithms and four regression algorithms that use diverse methodologies, including tree-, probability-, kernel-, ensemble-, and distance-based approaches. For both types of algorithm, we artificially subsampled the data to varying degrees, aiming to understand whether training models based on relatively extreme outcomes would yield improved performance. We also performed an information-gain analysis to examine which genes were most predictive of drug responses. Finally, we used tumor data from The Cancer Genome Atlas to evaluate the feasibility of predicting clinical responses in humans based on models derived from cell lines. When using classification or regression algorithms to predict discrete or continuous responses, respectively, we consistently observed excellent predictive performance when the training and test sets both consisted of cell-line data. However, classification models derived from cell-line data failed to generalize effectively for tumors.

## Introduction

Cancer is a complex and dynamic disease characterized by aberrant cellular processes such as excessive proliferation, resistance to apoptosis, and genomic instability (Hanahan and Weinberg, 2011). Tumors are caused by somatic variations, which can affect individual nucleotides or larger segments of DNA (Yao and Dai, 2014). Dysregulation of cellular function can also be caused by epigenetic modifications, including aberrant DNA methylation (Esteller et al., 2001). One goal of cancer research is to advance personalized medicine through identifying genomic and epigenomic features that influence treatment outcomes in individuals (McLeod, 2013). In this context, therapeutic decisions have the potential to be guided by molecular signatures.

Cancer cell lines are cell cultures developed from tumor samples. They represent one of the least expensive and most studied preclinical models (Masters, 2000). Drug screening in cell lines can be used to prioritize candidate drugs for testing in humans. In performing a screen, researchers calculate IC_50_ values, which quantify the amount of drug necessary to induce a biological response in half of the cells tested for a given experiment (Sebaugh, 2011). Drugs with a relatively high potency (corresponding to low log-transformed IC_50_ values) are generally considered to be the strongest candidates for use in humans, although patient safety must also be evaluated. After a candidate drug has been identified, researchers may seek to identify molecular markers associated with those responses, comparing cell lines that respond to the drug against those that do not. Such markers might be useful for elucidating drug mechanisms or eventually predicting clinical responses in patients (Iorio et al., 2016).

Over the past decade, researchers have catalogued the molecular profiles of more than a thousand cancer cell lines and their responses to hundreds of drugs (Barretina et al., 2012; Yang et al., 2013; Rees et al., 2016). These resources have been made publicly available, thus providing an opportunity for researchers to identify molecular signatures that predict drug responses in a preclinical setting. In addition, recent efforts to catalog molecular profiles in human tumors have resulted in massive collections of publicly available molecular data for tumor samples (ICGC, 2010; Tomczak et al., 2015; Forbes et al., 2017). Such data can be used to validate findings from preclinical studies and assess our ability to classify cancer patients into groups that will most likely benefit from a certain treatment (Azuaje, 2017).

Recent cell-line studies have emphasized the potential to predict drug responses based on gene-expression profiles (Costello et al., 2014; Yuan et al., 2014; Zhao et al., 2015; Chiu et al., 2019; Parca et al., 2019). Technologies for profiling gene-expression levels are widely available and reflect the downstream effects of genomic and epigenomic aberrations. However, gene-expression profiles may be difficult to apply in the clinic because of the instability of RNA (Geeleher et al., 2017). Moreover, gene-expression data are generated using a wide range of technologies (e.g., different types of oligonucleotide microarrays and RNA-sequencing), and are preprocessed using diverse algorithms. Thus, it may be difficult to combine datasets from multiple sources (e.g., preclinical and tumor data). In this study, we focus on DNA methylation profiles using cell-line data from the Genomics of Drug Sensitivity in Cancer (GDSC) database (Iorio et al., 2016) in combination with tumor data from The Cancer Genome Atlas (TCGA) (Hutter and Zenklusen, 2018). These projects used the same technology to quantify methylation levels and to perform data normalization, thus enabling us to perform a systematic evaluation of whether DNA methylation levels can predict drug responses.

DNA methylation is an epigenetic mechanism that controls gene-expression levels. The addition of a methyl group to DNA may lead to changes in DNA stability, chromatin structure, and DNA-protein interactions. Hypermethylation of CpG islands in promoter regions of DNA has been acknowledged as an important means of gene inactivation, and its occurrence has been detected in almost all types of human tumors (Esteller, 2002). Similar to genetic alterations, methylation changes to DNA may alter a gene’s behavior. However, hypermethylation can be reversed with the use of targeted therapy (Szyf, 2008), making it an attractive biomarker for anticancer therapy (Szyf, 1994; Arechederra et al., 2018).

In some cases, DNA methylation levels for a single gene may control cellular responses for a given drug. For example, MGMT hypermethylation predicts temozolomide responses in glioblastomas (Hegi et al., 2005), and BRCA1 hypermethylation predicts responses to poly ADP ribose polymerase inhibitors in breast carcinomas (Island, 2010). However, in many cases, drug responses are likely influenced by the combined effects of many genes interacting in the context of signaling pathways (Faivre et al., 2006). Accordingly, to maximize our ability to predict drug responses, it is critical to account for this complexity.

In this study, we use DNA methylation profiles from preclinical samples to model drug responses for eight anti-cancer drugs. We compare the performance of five classification algorithms and four regression algorithms that encompass a diverse range of methodologies, including tree-based, probability-based, kernel-based, ensemble-based, and distance-based approaches. For regression, we predict IC_50_ values directly. For classification, we use discretized IC_50_ values. For both types of algorithm, we artificially subsample the data to varying degrees to evaluate whether training models based on relatively extreme outcomes would yield improved performance; we assess our ability to predict drug responses using as few as 10% of the cell lines (those with the most extreme IC_50_ values). For each evaluation, we perform hyperparameter optimization of the algorithms. Perhaps surprisingly, classification algorithms performed best when only 10-20% of the cell lines were used. This outcome suggests that classification performance was most impacted by how extreme the IC_50_ values were, rather than by the amount of training data itself. In contrast, regression algorithms performed best when models were generated using all available cell lines. Finally, we derived classification models from the cell-line data and predicted drug responses for TCGA patients. However, in this setting, the models failed to generalize effectively, perhaps due to batch effects or other systematic biases.

## Methods

The GDSC database contains data for human cell lines derived from common and rare types of adult and childhood cancers. GDSC provides multiple types of molecular data for these cell lines in addition to response values for 265 anti-cancer drugs. In this work, we use database version GDSC1, which includes data for 987 cell lines curated between 2010 to 2015 (Iorio et al., 2016). Drug responses were measured as the natural log of the fitted IC_50_ value, which indicates the drug concentration required to induce a 50% reduction in cell proliferation *in vitro*. The more sensitive the cell line, the lower the IC_50_ value will be for any given drug. We developed machine-learning models of drug response using DNA methylation data from GDSC1 that had been preprocessed and summarized as gene-level *beta* values (Iorio et al., 2016); these values range between 0 and 1 (higher values indicate relatively high methylation for a given gene).

For external validation, we used DNA methylation and clinical drug-response data from patients in TCGA. We selected eight drugs that were administered to TCGA patients and were also present in the GDSC database: Gefitinib, Cisplatin, Docetaxel, Doxorubicin, Etoposide, Gemcitabine, Paclitaxel, and Temozolomide. These drugs represent a variety of molecular mechanisms, including DNA crosslinking, microtubule stabilization, and pyrimidine anti-metabolization. Aside from Gefitinib, which we used for model optimization on GDSC only, these drugs were associated with the largest number of patient drug-response values in TCGA (Huang et al., 2020). The GDSC database provides DNA methylation values for 6,035 TCGA samples that have been preprocessed using the same pipeline as the GDSC samples. We obtained drug-response data for TCGA patients from (Ding et al., 2016).

Cell lines with missing IC_50_ values were excluded on a per-drug basis; thus, sample sizes differed across the drugs. We started with a classification analysis, which aims to distinguish samples belonging to two or more pre-defined groups. Classification algorithms are widely available, and their predictions are intuitive to interpret—they assign probabilities to each sample for each class. To enable classification for the GDSC cell lines, we discretized the IC_50_ values into “low” and “high” values. However, the choice of a threshold for distinguishing low and high values was necessarily arbitrary. Initially, we used the median IC_50_ value across all cell lines as a threshold. However, cell lines with an IC_50_ just above or below this threshold naturally showed very little difference in their drug responses, even though they were assigned to different classes. In contrast, cell lines with extreme IC_50_ values (far from the threshold) had much more distinct drug responses. To investigate the effect of using a threshold to discretize the IC_50_ values for classification, we used subsampling. We created 10 different scenarios that included increasing percentages of the overall data. First, we sorted the data by IC_50_ values in ascending order. For the first scenario, we evaluated cell lines with the 5% lowest and 5% highest IC_50_ values (10% of the total data). In the next scenario, we evaluated cell lines with the 10% lowest and 10% highest IC_50_ values (20% of the total data), and so on. The last scenario included all the data, where the lowest 50% were considered to have low IC_50_ values and the highest 50% were considered to have high values (Figure S1). For the regression analysis, we followed a similar process for subsampling but retained the continuous nature of the IC_50_ values.

For both classification and regression, we used Random Forests (tree-based) (Breiman, 2001), Support Vector Machines (kernel-based) (Vapnik, 1998), Gradient Boosting Machines (ensemble-based) (Breiman, 1997), and k-nearest Neighbors (distance-based) (Cover and Hart, 1967) algorithms. The Naïve Bayes (probability-based) (Maron, 1961) algorithm was also used for classification but not for regression; this algorithm is only designed for classification analysis. We performed the analyses using the R programming language (R Core Team, 2019) and Rstudio (https://rstudio.com). The machine-learning algorithms are implemented in the following R packages: mlr (Bischl et al., 2016), e1071 (Meyer et al., 2019), xgboost (Chen et al., 2015), randomForest (Liaw and Wiener, 2002), and kknn (Schliep and Hechenbichler, 2016).

Using the GDSC cell-line data, we sought to select the best hyperparameters for each algorithm via nested cross validation. We used the *mlr* package (Bischl et al., 2016) to randomly assign the cell lines to 10 outer folds and 5 inner folds (per outer fold). For each combination of algorithm and data-subsampling scenario, we evaluated the performance of all hyperparameter combinations (Table 1) using the inner folds; we used MMCE (Mean Misclassification Error) (Schiffner et al., 2016) for classification and MSE (Mean Squared Error) (Hyndman, 2006) for regression as evaluation metrics (defaults in *mlr*). For the outer-fold predictions, we assessed performance for predicting drug response using several performance metrics. This enabled us to evaluate how consistently the algorithms performed. For the classification analysis, we used accuracy (1 - MMCE), area under the receiver operating characteristic curve (AUC) (Fan et al., 2006), F1 measure (Forman, 2003), Matthews correlation coefficient (MCC) (Baldi et al., 2000), recall, and specificity. For the regression analysis, we used MSE, Mean Absolute Error (MAE), and Root Mean Square Error (RMSE) (Hyndman, 2006).

**Table 1:**
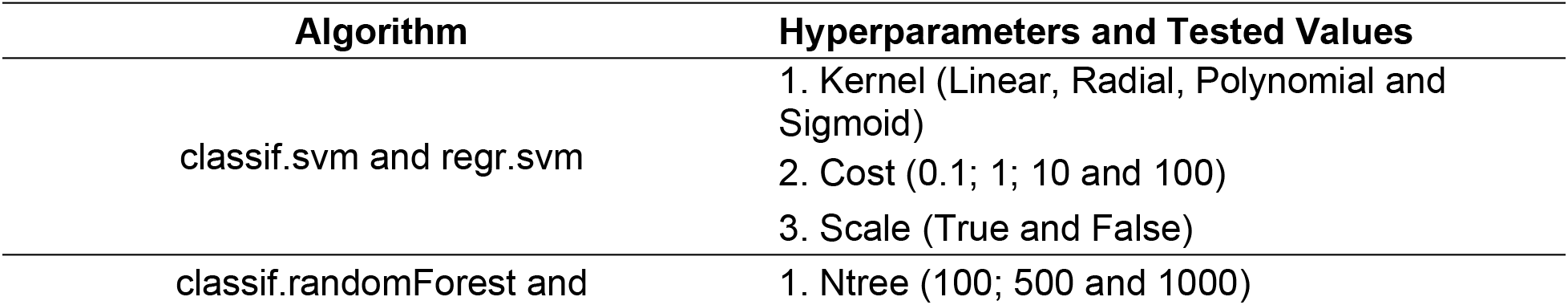
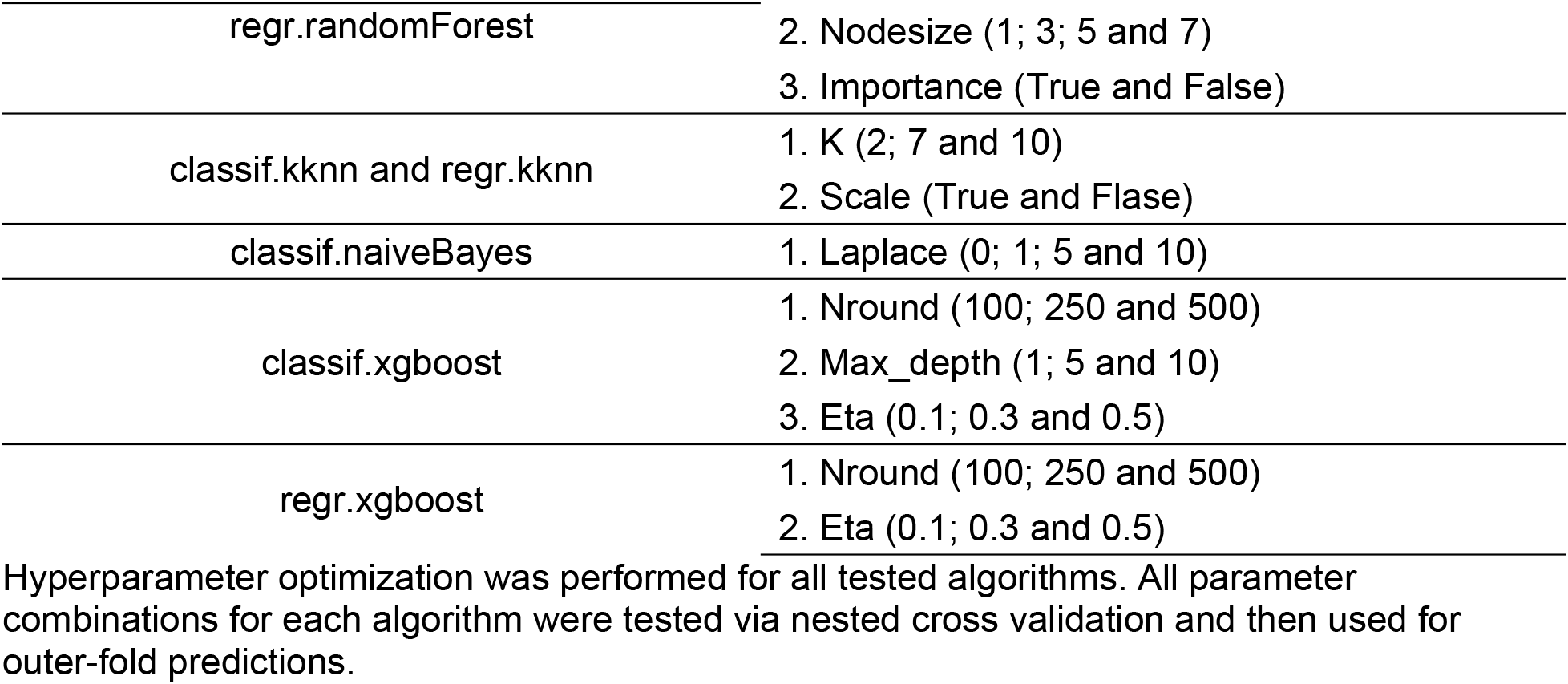
Hyperparameters Evaluated.

For the classification and regression analyses, we used feature selection to identify genes deemed to be most informative. We performed an information-gain analysis, assigning an importance score to each feature (gene). This analysis was implemented using the FSelectorRcpp package (Zawadzki and Kosinski, 2020). To assess the functional relevance of the top-ranked genes, we used the Panther Classification System to perform a statistical overrepresentation test based on protein classes (Mi et al., 2013). We used Fisher’s exact test to assess statistical significance and the False Discovery Rate to adjust for multiple testing (Benjamini and Hochberg, 1995).

For additional validation, we trained models based on drug responses in the GDSC cell lines and then predicted patient drug responses using tumor data from TCGA. These patient responses were based on clinical data, having no direct relation to IC50 values. Because the patient-response values were categorical in nature, we only performed classification for these data. We performed hyperparameter optimization using the GDSC (training) data.

## Results

Using data from 987 cell lines, we used machine-learning algorithms to evaluate the potential to predict cytotoxic responses based on genome-wide, DNA methylation profiles. Second, we examined which genes were most predictive of these responses. Finally, we evaluated the feasibility of predicting clinical responses in humans based on models derived from cell-line data.

### Classification analysis using cell-line data

We collected DNA methylation data and IC_50_ response values for eight drugs from the GDSC repository. In our initial analysis, we aimed to predict categories (classes) of drug sensitivity. These categories represented each cell line’s “response” or “non-response” to the drugs, corresponding to relatively low or high IC_50_ values, respectively. This categorization facilitated a simplified yet intuitive interpretation of the treatment outcomes and enabled us to use classification algorithms, which have been implemented for a broader range of algorithmic methodologies than regression algorithms.

Before performing classification, we categorized each cell line on a per-drug basis according to whether its IC_50_ value was greater than the median across all cell lines. One limitation of categorizing the cell lines in this way is that cell lines just above or below the median threshold show a relatively small difference in IC_50_ values, even though they are assigned to different classes. Generally, IC_50_ values did not follow a multimodal distribution (Figure 1). Therefore, we evaluated whether classification performance could be improved by excluding cell lines with an IC_50_ value relatively close to the median, even though this would reduce the amount of data available for training and testing. We evaluated ten scenarios that varied the number of cell lines used. In the most extreme scenario, we used methylation data for the cell lines with the 5% lowest and 5% highest IC_50_ values. In describing these subsampling scenarios, we use a notation that indicates the percentage of samples on each side of the distribution as well as the algorithm type. For example, when we analyzed the samples with the 5% highest and 5% lowest IC_50_ values and employed a classification algorithm, we indicate this using +-5%c. The equivalent scenario for regression would be represented as +-5%r.

**Figure 1:**
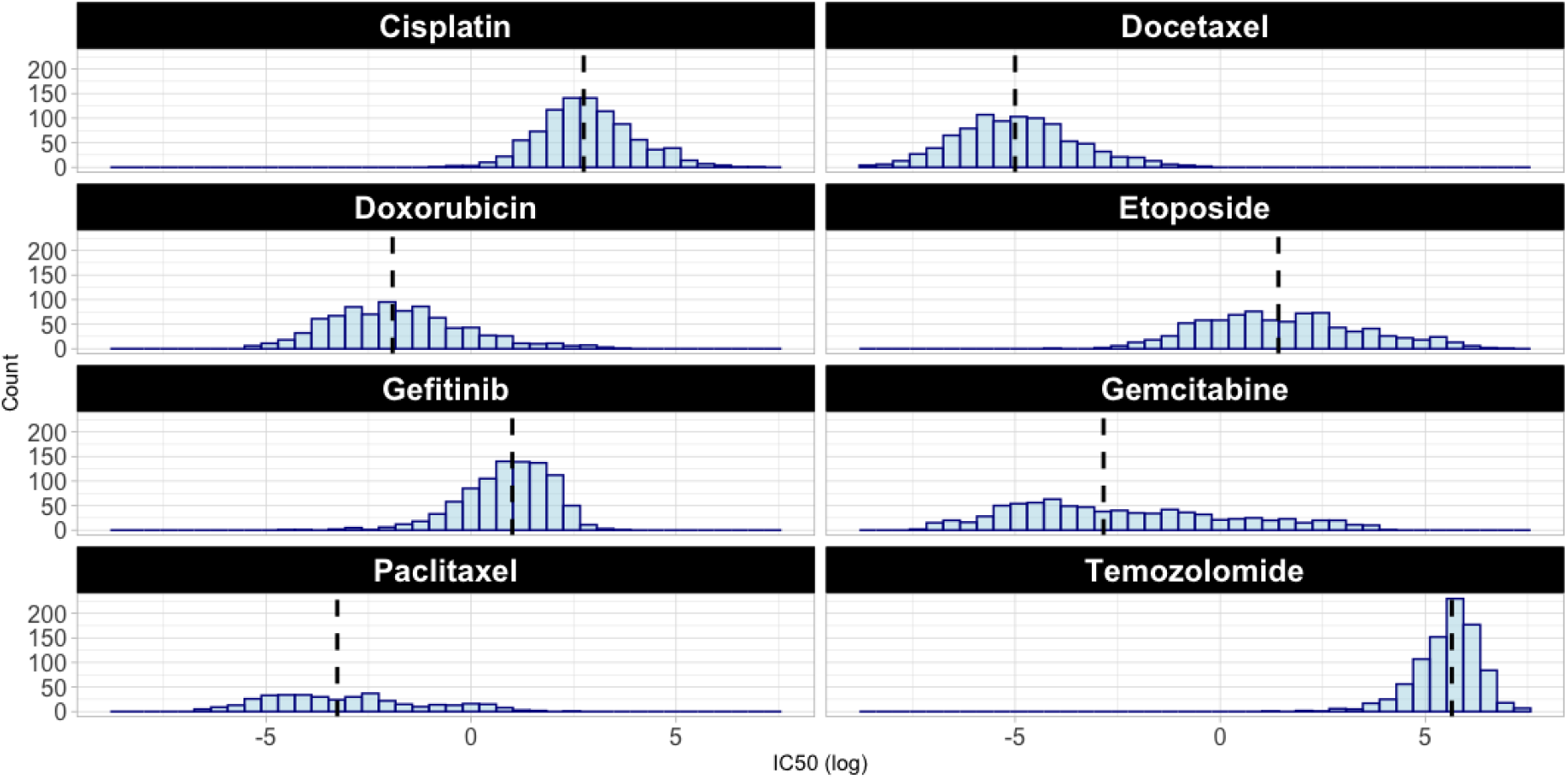
Histograms for each drug based on drug response (IC_50_ values) from the GDSC dataset. The black line represents the median value for each subsample across all available cell lines for each drug.

For this analysis, we applied five classification algorithms and evaluated their performance using nested cross validation and six performance metrics (see Methods). Initially, we evaluated Gefitinib, an EGFR inhibitor. The classification algorithms performed best when relatively few cell lines (those with the most extreme IC_50_ values) were used to train and test the models, attaining an AUC value as high as 0.94 (Table 2). Overall best performance was obtained for +-10%c or +-20%c. This pattern was consistent across all five algorithms and all six metrics that we evaluated (Figure 2). However, the Support Vector Machines (SVM) algorithm consistently achieved higher classification performance than the other algorithms for this drug.

**Table 2:**
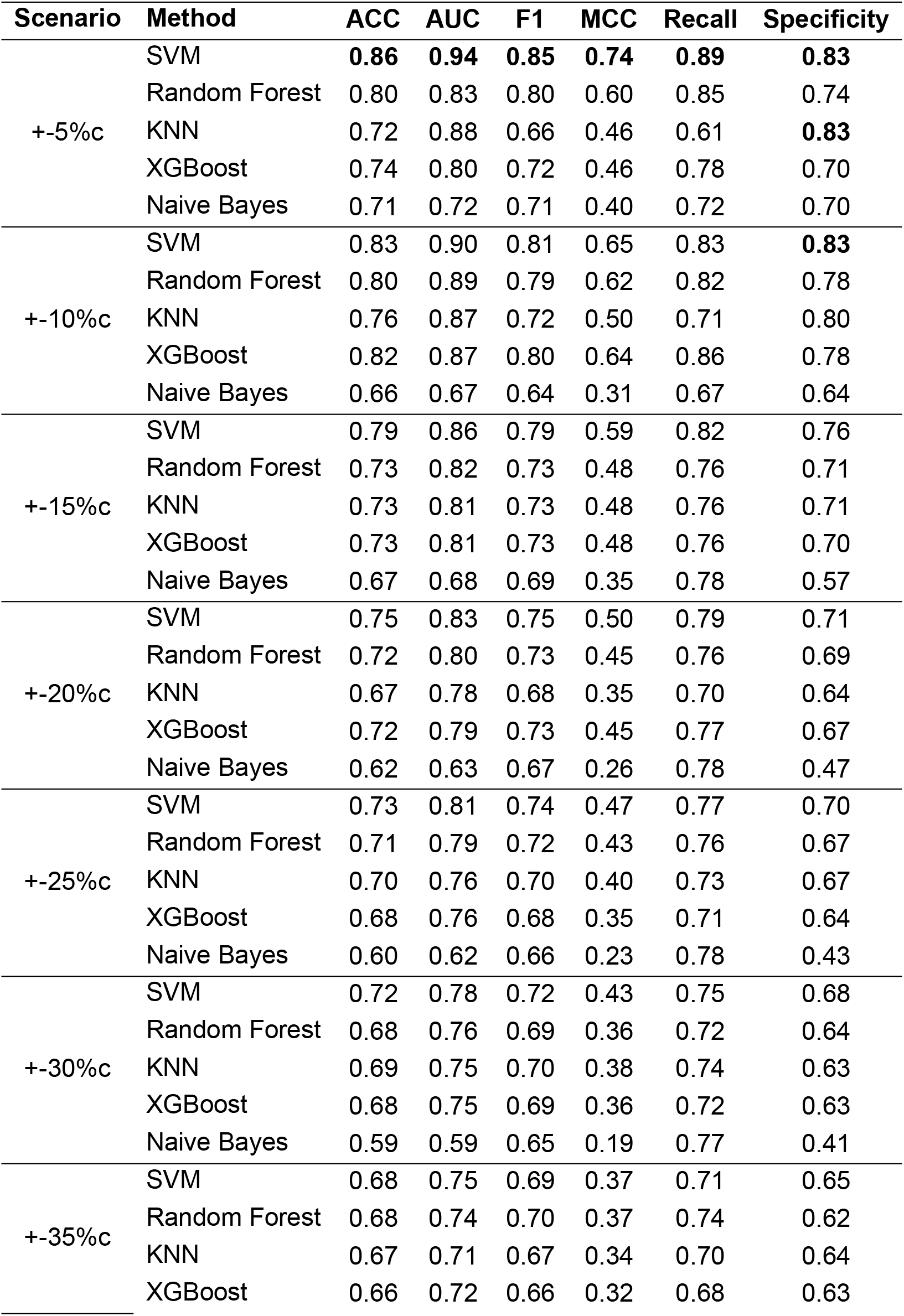
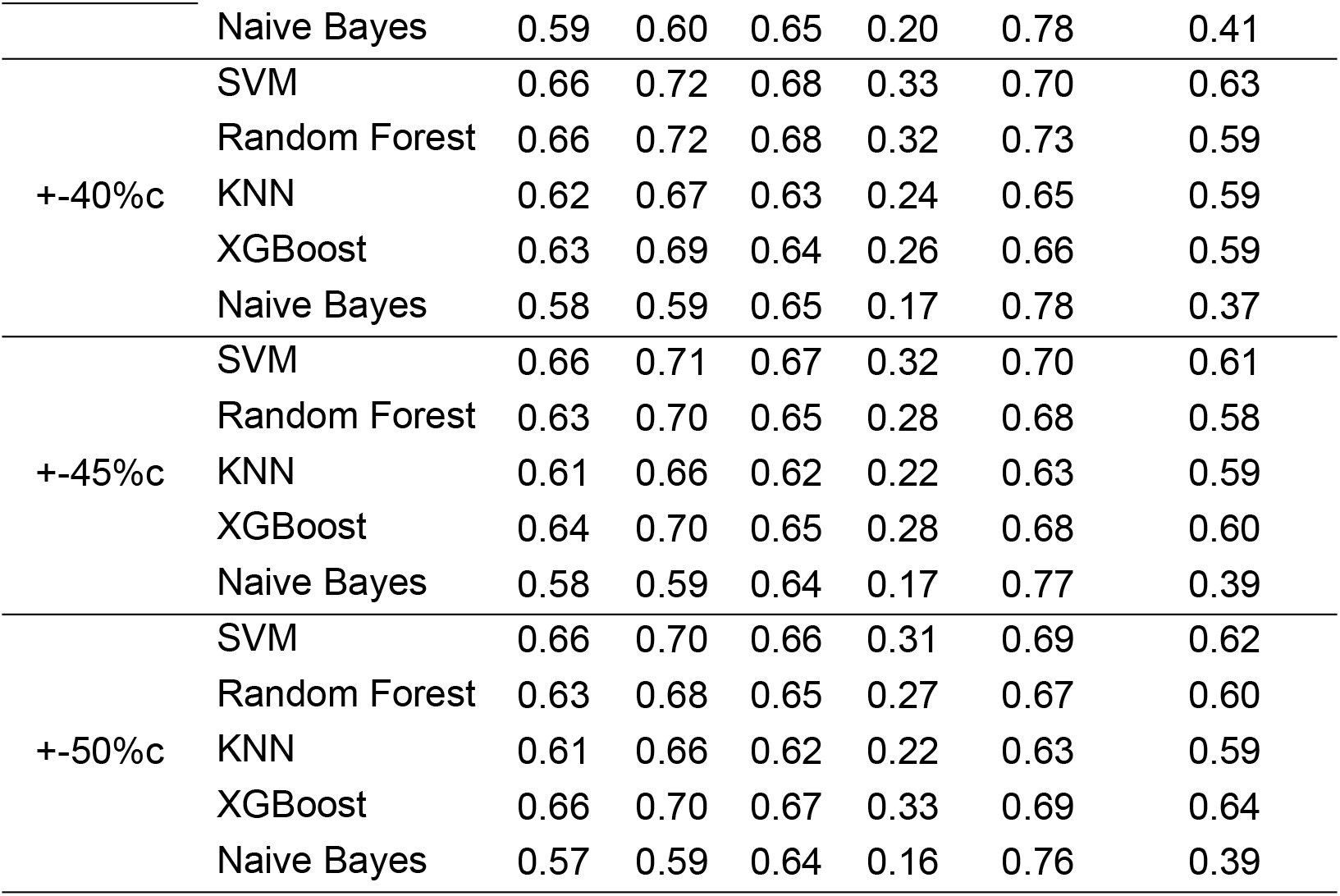
Classification results for all combinations of subsampling scenarios and algorithms for Gefitinib.

**Figure 2:**
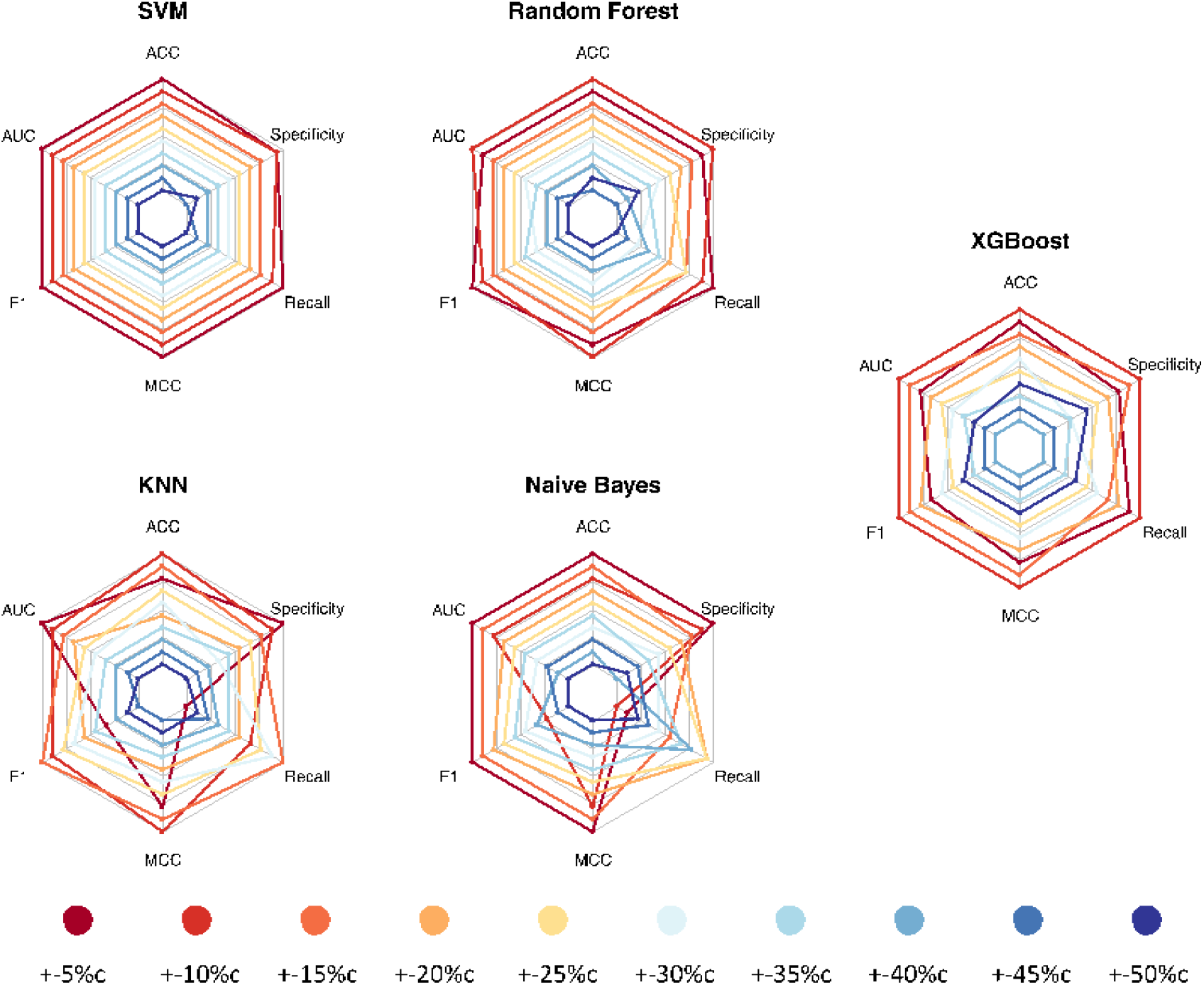
Graphs for Gefitinib classification analysis. The graphs compare different scenarios ranked in order of best result. GDSC cell-line data was used to generate ten subsampling scenarios, which we then tested via nested cross validation. Scenarios that are further away from the center represent higher metric values than scenarios closer to it. The evaluated metrics for each algorithm are accuracy (ACC), specificity, recall, Matthews correlation coefficient (MCC), F1 score (F1) and area under the receiver operating characteristic curve (AUC). Scenarios that used relatively few cell lines—but those with the most extreme IC_50_ values— performed best for all algorithms. Specific metric values may be found in Table 2.

For the seven remaining drugs, we continued to see a trend in which using a relatively small proportion of the data resulted in better classification performance. For Cisplatin, Docetaxel, Doxorubicin, and Etoposide, best performance was attained for +-5%c and +-10%c, and the best-performing algorithms were always SVM, Random Forests (RF), or k-nearest neighbors (KNN) (Tables S1-S7). In contrast, for Gemcitabine, the highest AUC value (0.80) was obtained for +-15%c (SVM algorithm). For Paclitaxel, the Gradient Boosting Machines (XGBoost) algorithm performed best for +-35%c (AUC = 0.75). The overall highest AUC value was attained for Docetaxel (0.96, +-10%c, Random Forests). Figures S2-S8 illustrate these results across all algorithms, metrics, and drugs and show that generally the top-performing algorithms were consistent across all the metrics, although these patterns were less consistent in scenarios where the highest AUC values were lower than 0.80.

To further analyze combinations of subsampling scenarios and classification algorithms, we ranked the AUC values for all combinations and each drug (where the lowest rank was considered best and represented the highest AUC value). Subsequently, we calculated the average AUC rank across all drugs (a lower AUC rank average indicated a better result). The best performance was attained for +-5%c SVM and +-10%c RF, achieving average ranks of 4.29 and 6.38, respectively (Table 3). Evaluating the minimum, mean and maximum AUC values for each combination of drug and algorithm, Docetaxel attained the best overall performance based on AUC (Table 4).

**Table 3:** Average AUC rank for all combinations of subsampling scenarios and algorithms.

| Scenario | Method | Average AUC Rank |
| --- | --- | --- |
| +-5%c | SVM | 4.29 |
| +-10%c | Random Forest | 6.38 |
| +-5%c | Random Forest | 6.71 |
| +-10%c | SVM | 8.00 |
| +-10%c | XGBoost | 9.38 |
| +-15%c | SVM | 9.50 |
| +-15%c | Random Forest | 9.75 |
| +-20%c | SVM | 10.50 |
| +-15%c | XGBoost | 10.88 |
| +-10%c | KNN | 12.13 |
| +-5%c | KNN | 12.71 |
| +-25%c | SVM | 13.00 |
| +-25%c | XGBoost | 13.75 |
| +-5%c | XGBoost | 14.00 |
| +-25%c | Random Forest | 15.50 |
| +-30%c | SVM | 15.88 |
| +-20%c | XGBoost | 16.38 |
| +-20%c | Random Forest | 16.50 |
| +-30%c | XGBoost | 17.25 |
| +-35%c | SVM | 19.88 |
| +-35%c | XGBoost | 20.38 |
| +30%<sub>c</sub> | Random Forest | 21.13 |
| +15%<sub>c</sub> | KNN | 21.50 |
| +25%<sub>c</sub> | KNN | 25.00 |
| +20%<sub>c</sub> | KNN | 25.50 |
| +30%<sub>c</sub> | KNN | 25.63 |
| +35%<sub>c</sub> | Random Forest | 26.13 |
| +40%<sub>c</sub> | SVM | 26.25 |
| +40%<sub>c</sub> | Random Forest | 27.50 |
| +45%<sub>c</sub> | SVM | 27.50 |
| +40%<sub>c</sub> | XGBoost | 28.25 |
| +5%<sub>c</sub> | Naive Bayes | 28.57 |
| +45%<sub>c</sub> | XGBoost | 30.13 |
| +50%<sub>c</sub> | SVM | 31.00 |
| +35%<sub>c</sub> | KNN | 32.25 |
| +45%<sub>c</sub> | Random Forest | 32.25 |
| +50%<sub>c</sub> | XGBoost | 32.88 |
| +10%<sub>c</sub> | Naive Bayes | 34.38 |
| +40%<sub>c</sub> | KNN | 36.63 |
| +50%<sub>c</sub> | Random Forest | 37.75 |
| +15%<sub>c</sub> | Naive Bayes | 39.88 |
| +45%<sub>c</sub> | KNN | 40.00 |
| +50%<sub>c</sub> | KNN | 41.00 |
| +20%<sub>c</sub> | Naive Bayes | 42.88 |
| +25%<sub>c</sub> | Naive Bayes | 44.13 |
| +30%<sub>c</sub> | Naive Bayes | 44.38 |
| +35%<sub>c</sub> | Naive Bayes | 45.25 |
| +40%<sub>c</sub> | Naive Bayes | 46.50 |
| +45%<sub>c</sub> | Naive Bayes | 47.50 |
| +50%<sub>c</sub> | Naive Bayes | 48.88 |

AUC values were ranked for each drug and were then averaged. Lower ranks imply a better metric value.

**Table 4:** Minimum, mean and maximum AUC value for each drug and algorithm combination.

| **Drug** | **Method** | **Min** | **Mean** | **Max** |
| --- | --- | --- | --- | --- |
| Gefitinib | SVM | 0.70 | 0.80 | 0.94 |
| Gefitinib | Random Forest | 0.68 | 0.77 | 0.89 |
| Gefitinib | Naïve Bayes | 0.59 | 0.63 | 0.72 |
| Gefitinib | KNN | 0.66 | 0.75 | 0.88 |
| Gefitinib | XGBoost | 0.69 | 0.76 | 0.87 |
| Cisplatin | SVM | 0.68 | 0.78 | 0.90 |
| Cisplatin | Random Forest | 0.66 | 0.76 | 0.86 |
| Cisplatin | Naïve Bayes | 0.59 | 0.63 | 0.72 |
| Cisplatin | KNN | 0.60 | 0.72 | 0.84 |
| Cisplatin | XGBoost | 0.69 | 0.77 | 0.88 |
| Paclitaxel | SVM | 0.52 | 0.64 | 0.69 |
| Paclitaxel | Random Forest | 0.61 | 0.67 | 0.70 |
| Paclitaxel | Naïve Bayes | 0.53 | 0.57 | 0.60 |
| Paclitaxel | KNN | 0.58 | 0.65 | 0.70 |
| Paclitaxel | XGBoost | 0.58 | 0.68 | 0.75 |
| Temozolomide | SVM | 0.73 | 0.84 | 0.94 |
| Temozolomide | Random Forest | 0.72 | 0.82 | 0.89 |
| Temozolomide | Naïve Bayes | 0.63 | 0.68 | 0.75 |
| Temozolomide | KNN | 0.67 | 0.77 | 0.85 |
| Temozolomide | XGBoost | 0.73 | 0.82 | 0.92 |
| Etoposide | SVM | 0.66 | 0.75 | 0.89 |
| Etoposide | Random Forest | 0.62 | 0.70 | 0.85 |
| Etoposide | Naïve Bayes | 0.56 | 0.61 | 0.75 |
| Etoposide | KNN | 0.60 | 0.67 | 0.84 |
| Etoposide | XGBoost | 0.65 | 0.72 | 0.80 |
| Gemcitabine | SVM | 0.66 | 0.73 | 0.80 |
| Gemcitabine | Random Forest | 0.63 | 0.70 | 0.77 |
| Gemcitabine | Naïve Bayes | 0.56 | 0.59 | 0.74 |
| Gemcitabine | KNN | 0.60 | 0.66 | 0.75 |
| Gemcitabine | XGBoost | 0.62 | 0.71 | 0.79 |
| Docetaxel | SVM | 0.76 | 0.87 | 0.95 |
| Docetaxel | Random Forest | 0.76 | 0.86 | 0.96 |
| Docetaxel | Naïve Bayes | 0.65 | 0.71 | 0.79 |
| Docetaxel | KNN | 0.70 | 0.80 | 0.88 |
| Docetaxel | XGBoost | 0.76 | 0.86 | 0.95 |
| Doxorubicin | SVM | 0.64 | 0.70 | 0.78 |
| Doxorubicin | Random Forest | 0.59 | 0.67 | 0.80 |
| Doxorubicin | Naïve Bayes | 0.56 | 0.58 | 0.64 |
| Doxorubicin | KNN | 0.58 | 0.64 | 0.74 |
| Doxorubicin | XGBoost | 0.60 | 0.65 | 0.68 |

### Regression analysis using cell-line data

We performed a regression analysis using the same DNA methylation data and with non-discretized IC_50_ response values for the same eight drugs. For this analysis, we applied four regression algorithms and evaluated their performance using nested cross validation and three performance metrics (RMSE, MSE and MAE). As with the classification analysis, we performed data subsampling to evaluate the effects of using relatively extreme IC_50_ values. For Gefitinib, all algorithms performed best when all cell lines were used to train and test the models, attaining an RMSE value as low as 0.95 (lower is better, see Table 5). Similarly, for the other seven drugs, the best performance was attained when all data were used (Tables S8-S14). Across all drugs and metrics, the SVM and Random Forest algorithms performed best for every combination of drug and performance metric (Figure 3). Furthermore, predictive performance was highly consistent for all metrics used (Figures S9-S15). When evaluating the mean RMSE ranked values (where the lowest rank was considered best and represented the lowest RMSE value), the SVM and RF algorithms and the +-50%r scenarios performed best (Table 6), and predictions for Temozolomide were more accurate overall than those for other drugs (Table 7).

**Table 5:**
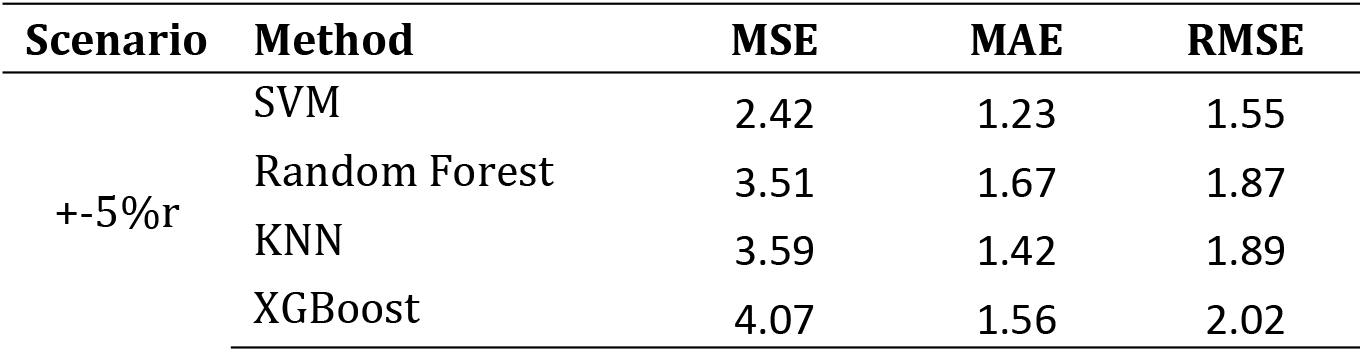
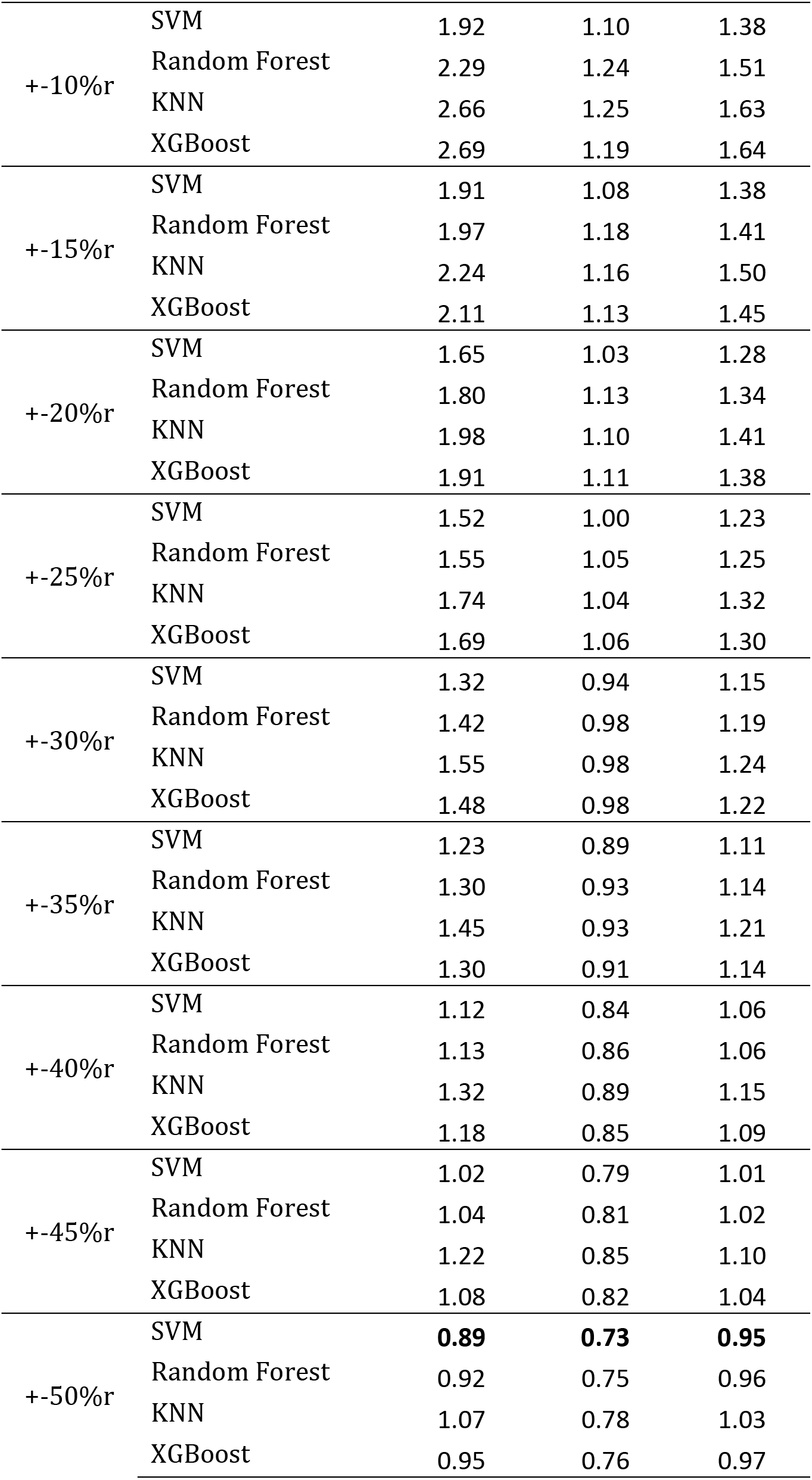
Regression results for all combinations of subsampling scenarios and algorithms for Gefitinib.

| Scenario | Method | MSE | MAE | RMSE |
| --- | --- | --- | --- | --- |
| +-5%r | SVM | 2.42 | 1.23 | 1.55 |
|  | Random Forest | 3.51 | 1.67 | 1.87 |
|  | KNN | 3.59 | 1.42 | 1.89 |
|  | XGBoost | 4.07 | 1.56 | 2.02 |
| +-10%r | SVM | 1.92 | 1.10 | 1.38 |
|  | Random Forest | 2.29 | 1.24 | 1.51 |
|  | KNN | 2.66 | 1.25 | 1.63 |
|  | XGBoost | 2.69 | 1.19 | 1.64 |
| +-15%r | SVM | 1.91 | 1.08 | 1.38 |
|  | Random Forest | 1.97 | 1.18 | 1.41 |
|  | KNN | 2.24 | 1.16 | 1.50 |
|  | XGBoost | 2.11 | 1.13 | 1.45 |
| +-20%r | SVM | 1.65 | 1.03 | 1.28 |
|  | Random Forest | 1.80 | 1.13 | 1.34 |
|  | KNN | 1.98 | 1.10 | 1.41 |
|  | XGBoost | 1.91 | 1.11 | 1.38 |
| +-25%r | SVM | 1.52 | 1.00 | 1.23 |
|  | Random Forest | 1.55 | 1.05 | 1.25 |
|  | KNN | 1.74 | 1.04 | 1.32 |
|  | XGBoost | 1.69 | 1.06 | 1.30 |
| +-30%r | SVM | 1.32 | 0.94 | 1.15 |
|  | Random Forest | 1.42 | 0.98 | 1.19 |
|  | KNN | 1.55 | 0.98 | 1.24 |
|  | XGBoost | 1.48 | 0.98 | 1.22 |
| +-35%r | SVM | 1.23 | 0.89 | 1.11 |
|  | Random Forest | 1.30 | 0.93 | 1.14 |
|  | KNN | 1.45 | 0.93 | 1.21 |
|  | XGBoost | 1.30 | 0.91 | 1.14 |
| +-40%r | SVM | 1.12 | 0.84 | 1.06 |
|  | Random Forest | 1.13 | 0.86 | 1.06 |
|  | KNN | 1.32 | 0.89 | 1.15 |
|  | XGBoost | 1.18 | 0.85 | 1.09 |
| +-45%r | SVM | 1.02 | 0.79 | 1.01 |
|  | Random Forest | 1.04 | 0.81 | 1.02 |
|  | KNN | 1.22 | 0.85 | 1.10 |
|  | XGBoost | 1.08 | 0.82 | 1.04 |
| +-50%r | SVM | **0.89** | **0.73** | **0.95** |
|  | Random Forest | 0.92 | 0.75 | 0.96 |
|  | KNN | 1.07 | 0.78 | 1.03 |
|  | XGBoost | 0.95 | 0.76 | 0.97 |

**Table 6:**
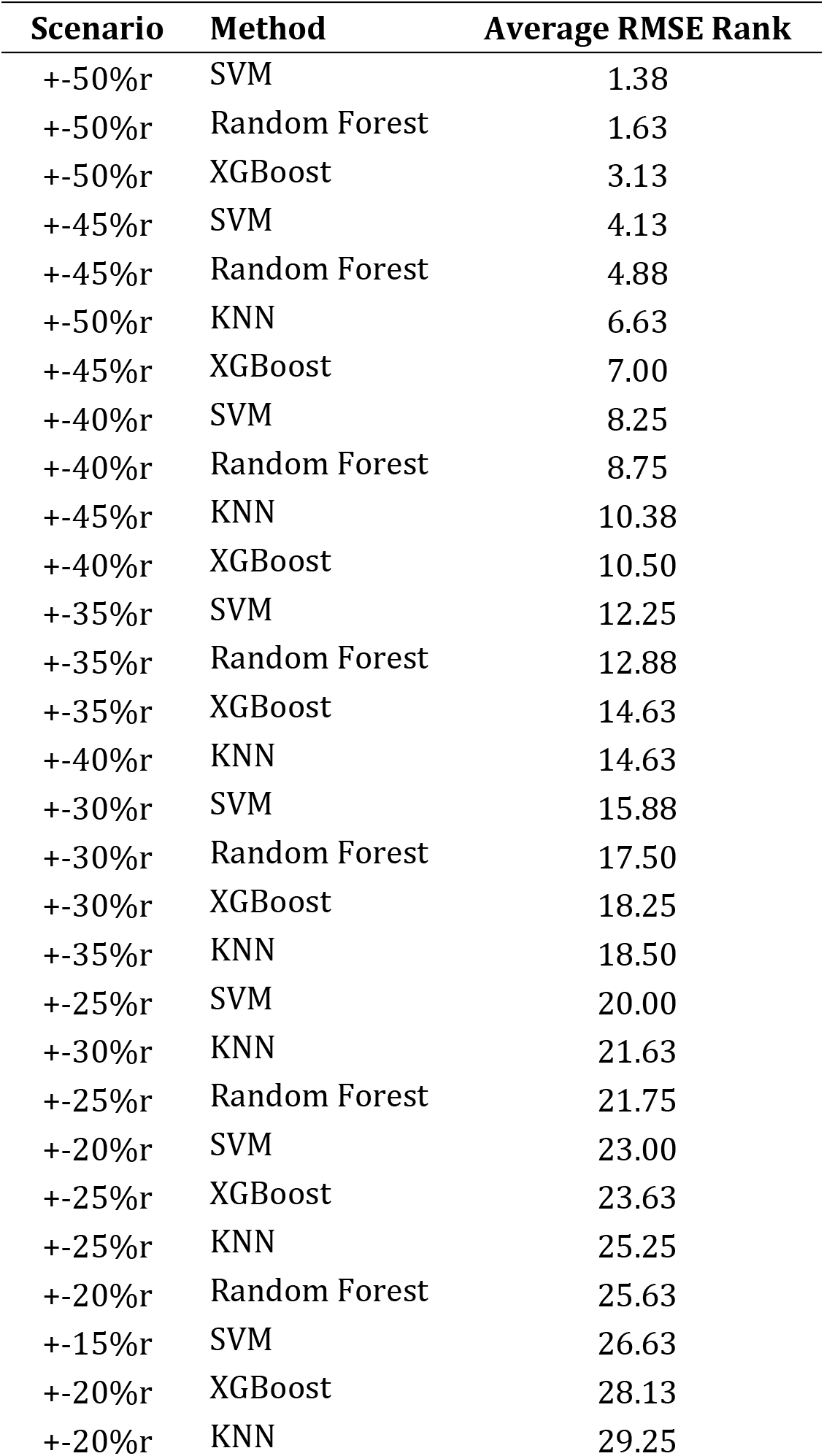
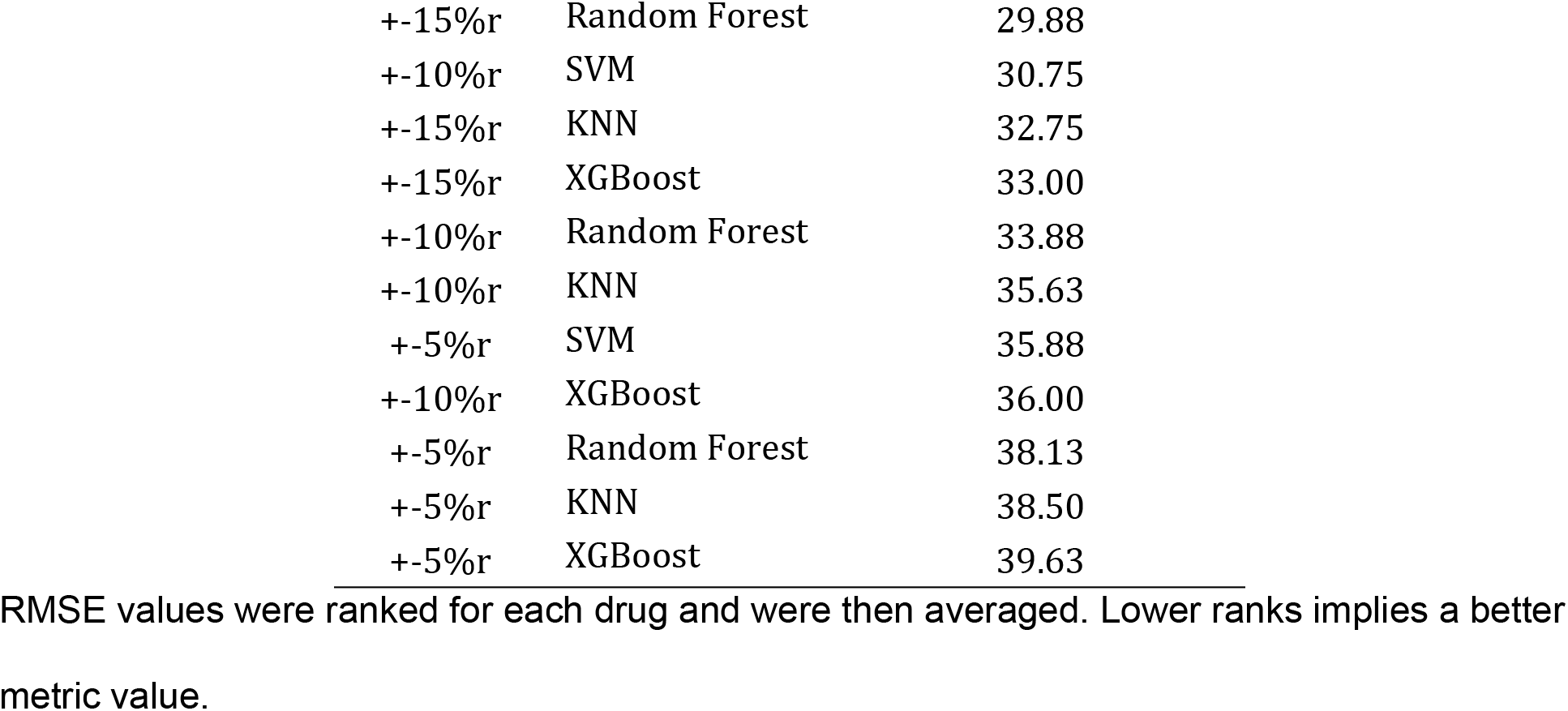
Average RMSE rank for all combinations of subsampling scenarios and algorithms.

| Scenario | Method | Average RMSE Rank |
| --- | --- | --- |
| +50%r | SVM | 1.38 |
| +50%r | Random Forest | 1.63 |
| +50%r | XGBoost | 3.13 |
| +45%r | SVM | 4.13 |
| +45%r | Random Forest | 4.88 |
| +50%r | KNN | 6.63 |
| +45%r | XGBoost | 7.00 |
| +40%r | SVM | 8.25 |
| +40%r | Random Forest | 8.75 |
| +45%r | KNN | 10.38 |
| +40%r | XGBoost | 10.50 |
| +35%r | SVM | 12.25 |
| +35%r | Random Forest | 12.88 |
| +35%r | XGBoost | 14.63 |
| +40%r | KNN | 14.63 |
| +30%r | SVM | 15.88 |
| +30%r | Random Forest | 17.50 |
| +30%r | XGBoost | 18.25 |
| +35%r | KNN | 18.50 |
| +25%r | SVM | 20.00 |
| +30%r | KNN | 21.63 |
| +25%r | Random Forest | 21.75 |
| +20%r | SVM | 23.00 |
| +25%r | XGBoost | 23.63 |
| +25%r | KNN | 25.25 |
| +20%r | Random Forest | 25.63 |
| +15%r | SVM | 26.63 |
| +20%r | XGBoost | 28.13 |
| +20%r | KNN | 29.25 |
| +15%r | Random Forest | 29.88 |
| +10%r | SVM | 30.75 |
| +15%r | KNN | 32.75 |
| +15%r | XGBoost | 33.00 |
| +10%r | Random Forest | 33.88 |
| +10%r | KNN | 35.63 |
| +5%r | SVM | 35.88 |
| +10%r | XGBoost | 36.00 |
| +5%r | Random Forest | 38.13 |
| +5%r | KNN | 38.50 |
| +5%r | XGBoost | 39.63 |

RMSE values were ranked for each drug and were then averaged. Lower ranks implies a better metric value.

**Table 7:** Minimum, mean and maximum RMSE value for each drug and algorithm combination.

| Drug | Method | Min | Mean | Max |
| --- | --- | --- | --- | --- |
| Gefitinib | SVM | 0.95 | 1.21 | 1.55 |
| Gefitinib | Random Forest | 0.96 | 1.27 | 1.87 |
| Gefitinib | KNN | 1.03 | 1.35 | 1.89 |
| Gefitinib | XGBoost | 0.97 | 1.32 | 2.02 |
| Cisplatin | SVM | 1.04 | 1.36 | 2.04 |
| Cisplatin | Random Forest | 1.03 | 1.37 | 2.11 |
| Cisplatin | KNN | 1.10 | 1.44 | 2.18 |
| Cisplatin | XGBoost | 1.05 | 1.42 | 2.06 |
| Paclitaxel | SVM | 1.90 | 2.52 | 3.52 |
| Paclitaxel | Random Forest | 1.86 | 2.52 | 3.63 |
| Paclitaxel | KNN | 1.97 | 2.64 | 3.67 |
| Paclitaxel | XGBoost | 1.91 | 2.74 | 4.54 |
| Temozolomide | SVM | 0.67 | 0.82 | 1.15 |
| Temozolomide | Random Forest | 0.67 | 0.87 | 1.31 |
| Temozolomide | KNN | 0.73 | 0.95 | 1.38 |
| Temozolomide | XGBoost | 0.70 | 0.91 | 1.56 |
| Etoposide | SVM | 1.79 | 2.32 | 2.95 |
| Etoposide | Random Forest | 1.84 | 2.41 | 3.06 |
| Etoposide | KNN | 1.94 | 2.50 | 3.07 |
| Etoposide | XGBoost | 1.87 | 2.57 | 3.80 |
| Gemcitabine | SVM | 2.60 | 3.26 | 4.13 |
| Gemcitabine | Random Forest | 2.55 | 3.36 | 4.52 |
| Gemcitabine | KNN | 2.70 | 3.53 | 4.50 |
| Gemcitabine | XGBoost | 2.63 | 3.48 | 4.79 |
| Docetaxel | SVM | 1.22 | 1.48 | 2.08 |
| Docetaxel | Random Forest | 1.24 | 1.54 | 2.21 |
| Docetaxel | KNN | 1.32 | 1.71 | 2.72 |
| Docetaxel | XGBoost | 1.24 | 1.60 | 2.44 |
| Doxorubicin | SVM | 1.57 | 2.15 | 3.22 |
| Doxorubicin | Random Forest | 1.58 | 2.19 | 3.29 |
| Doxorubicin | KNN | 1.70 | 2.24 | 3.10 |
| Doxorubicin | XGBoost | 1.60 | 2.27 | 3.51 |

**Figure 3:**
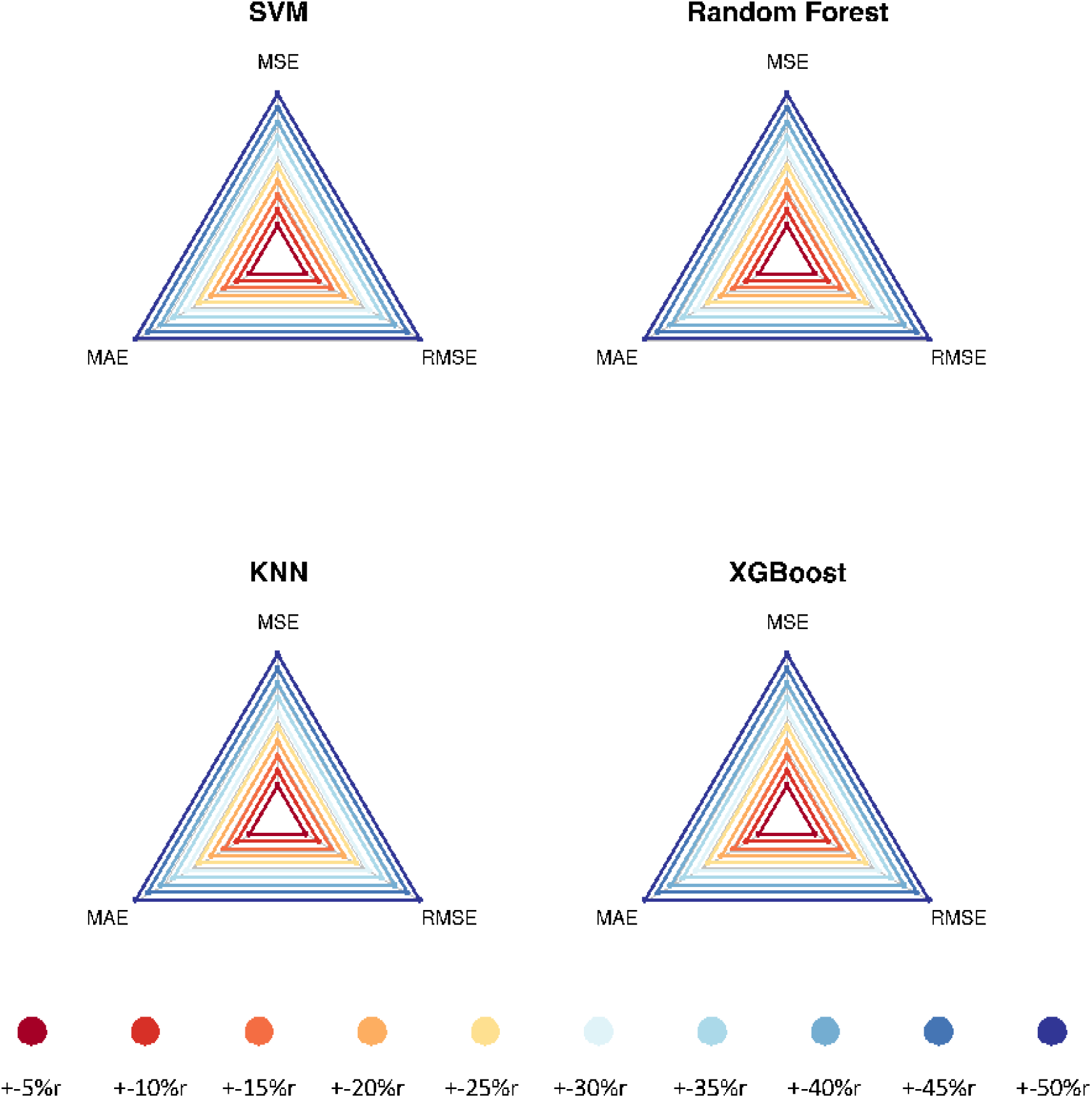
Graphs for Gefitinib regression analysis. We used DNA methylation data from cell lines to predict continuous IC_50_ response values using four regression algorithms. We evaluated the algorithms’ performance via nested cross validation for ten subsampling scenarios. The graphs illustrate the performance for these scenarios, ranked in order of relative performance for three metrics: MSE (Mean Squared Error), RMSE (Root Mean Square Error), and MAE (Mean Absolute Error). Scenarios further away from the center represent relatively low metric values (and thus better performance). Scenarios that used all cell lines performed best for all algorithms. Specific metric values may be found in Table 5.

### Informative genes for predicting cell-line responses

The DNA methylation assays target CpG islands associated with genes across the genome. After identifying analysis scenarios that resulted in optimal performance for classification and regression, we used feature selection to identify genes that were most informative for these scenarios. For the classification analysis, we focused on the +-5%c scenario. For the regression task, we focused on the +-50%r scenario. Table 8 lists the 20 top-ranked genes for Gefitinib. The *CTGF* gene was ranked 1st for the classification analysis and 15th for the regression analysis. The *CTGF* protein plays important roles in signaling pathways that control tissue remodeling via cellular adhesion, extracellular matrix deposition, and myofibroblast activation (Lipson, 2012); these processes are known to influence tumorigenesis and may alter drug responses (Hirohashi and Kanai, 2003). For example, EGFR is expressed in many head and neck squamous cell carcinomas and non-small cell lung carcinomas, yet many of these patients do not respond to Gefitinib treatment (Frederick et al., 2007). This lack of response has been associated with a loss of cell-cell adhesion, elongation of cells, and tumor-cell invasion of the extracellular matrix (Yauch et al., 2005; Thomson et al., 2005; Witta et al., 2006). F11R was ranked second in importance for the classification analysis and fourth for the regression analysis. The protein encoded by this gene is a junctional adhesion molecule that regulates the integrity of tight junctions and permeability (Naik and Eckfeld, 2003). Although these associations provide some support for our feature-selection results and that adhesion processes are important to Gefitinib responses, none of the other top-20 genes overlapped between the classification and regression analysis, and none of the other genes are known to be involved in cancer-related pathways (Kanehisa and Goto, 2000). However, we note that SVM and RF models represent multivariate patterns; thus, known cancer genes may alter drug responses in combination with the genes identified via our univariate feature-selection approach, even if they are not among the top-ranked genes.

**Table 8:**
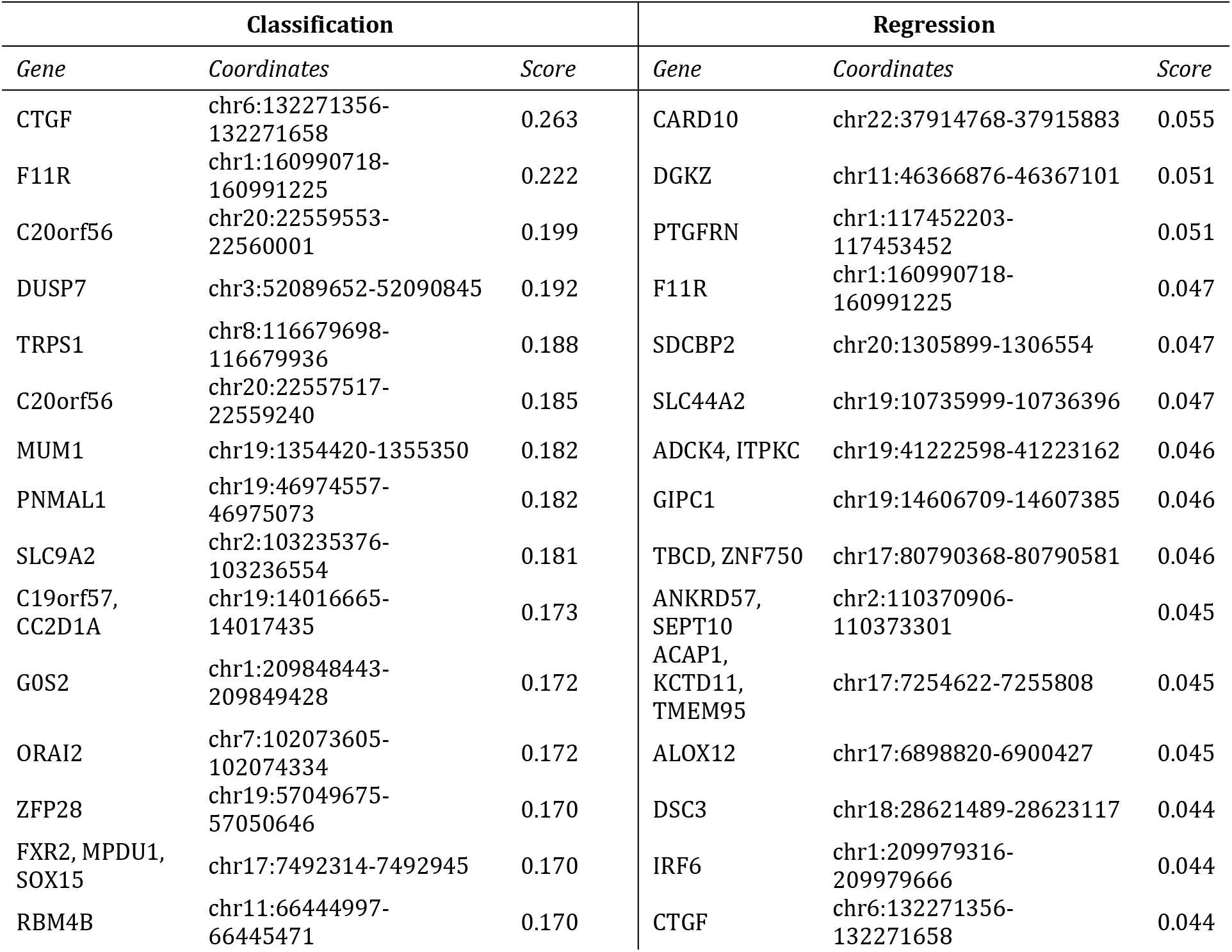
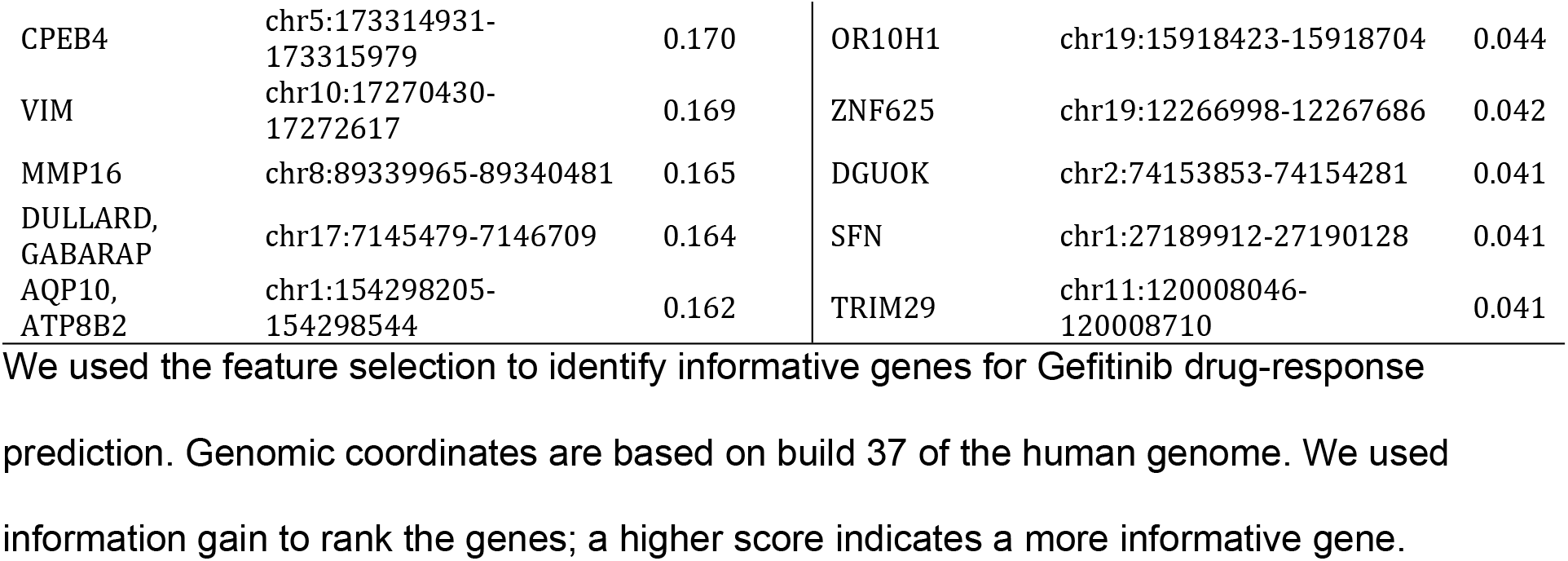
Informative genes for predicting cell-line responses for Gefitinib.

Tables S15-S21 provide the top-20 gene lists for the other 7 drugs. To gain insight regarding the roles that these genes might play in drug responses, we used the Panther Classification System to identify overrepresented protein classes associated with these genes. For the classification analysis, *tight junction* and *cell junction protein* were the only protein classes that resulted in false discovery rates lower than 0.25 (for Cisplatin and Doxorubicin) (Table S22). Similar results occurred for the regression analysis, but the *nucleotidyltransferase* protein class was significant for Doxorubicin, and the *kinase* and *cadherin* protein classes were significant for Gefitinib (Table S23).

### Using methylation profiles from cell lines to predict tumor/patient drug responses

The above analyses used methylation profiles to predict drug responses in cell lines. Via cross validation, we showed that high levels of predictive accuracy are attainable using this approach. We also found that smaller data sets with more extreme IC_50_ values yielded the best classification results and that the SVM and RF algorithms typically produced the most accurate results. Next we evaluated whether this performance would hold true in a translational-medicine context. The GDSC repository provides methylation profiles for 6,035 tumors from TCGA; these data have been preprocessed using the same methodology as the GDSC samples, thus enabling easier integration and reducing technical biases. For 1,638 TCGA patients, clinical drug-response information was available. These data indicate available outcomes over the course of the patients’ treatment by physicians (not as part of clinical trials). In many cases, drug-response values for multiple drugs were recorded for a given patient. Each response value was categorized as “clinical progressive disease,” “stable disease,” “partial response,” or “complete response”. These respective categories represent increasing levels of response to a given drug.

We trained the SVM and RF classification algorithms on the full GDSC dataset and predicted drug-response categories for each TCGA patient for which methylation and drug-response data were available. Based on our cross-validation results from the GDSC analysis, we focused on the +-5%c and +-10%c scenarios. For each TCGA test sample, our models generated a probabilistic prediction indicating whether that patient would respond to a given drug. We hypothesized that these probabilities would be associated with actual clinical responses in the patients. For example, we expected that the probabilities would be higher for partial and complete responders than for patients whose responses had been classified as clinical progressive disease or stable disease. To quantify this association, we used linear regression to calculate a slope representing our ability to predict clinical responses based on the classification probabilities. However, the slopes were consistently near zero (Table 9 and Figure 4), suggesting that the classification algorithms were unable to identify patterns in the cell-line methylation data that translate to patient responses. One exception was the RF predictions for Gefitinib response, but TCGA data were available only for 2 patients who had been treated with this drug.

**Table 9:** Slope values for selected combinations of subsampling scenarios and algorithms for all drugs.

| Drug | +5%c SVM | +5%c Random Forest | +10%c SVM | +10%c Random Forest |
| --- | --- | --- | --- | --- |
| Cisplatin (n=211) | -0.0091 | -0.0021 | -0.0059 | -0.0002 |
| Docetaxel (n=71) | 0.0342 | 0.0069 | 0.0418 | 0.0190 |
| Doxorubicin (n=79) | -0.0271 | -0.0166 | 0.0041 | -0.0015 |
| Etoposide (n=33) | 0.0304 | 0.0239 | 0.0088 | 0.0121 |
| Gefitinib (n=2) | 0.0103 | 0.0199 | 0.0181 | 0.2472 |
| Gemcitabine (n=102) | -0.0178 | -0.0079 | -0.0244 | -0.0145 |
| Paclitaxel (n=139) | 0.0050 | 0.0017 | 0.0030 | 0.0071 |
| Temozolomide (n=86) | -0.0064 | -0.0086 | 0.0034 | 0.0023 |

Slopes close to zero show that the evaluated models were unable to identify patterns that successfully translated into patient responses.

**Figure 4:**
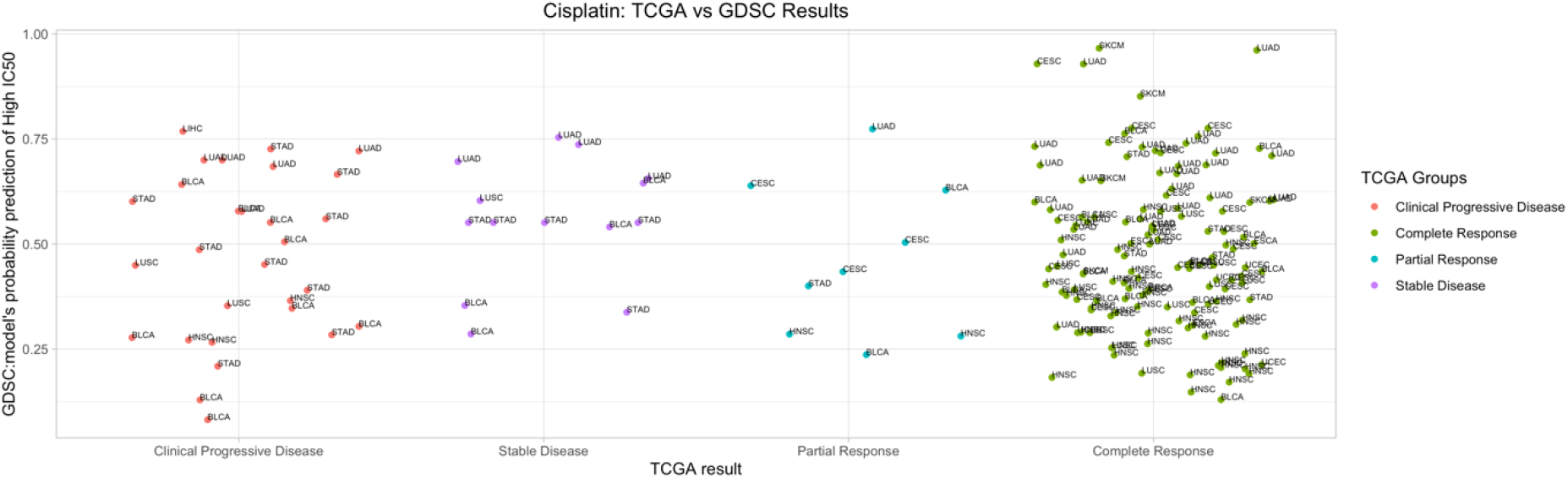
Predicting patient drug response from methylation profiles for Cisplatin (n=211). For each TCGA test sample, our model (+-10%c SVM) generated a probabilistic prediction to drug response. Our original hypothesis was that these probabilities would be associated with the patient’s clinical response. However, the classification algorithm was not able to identify active patterns to translate methylation profile tumor drug response prediction.

To this point, all of our classification models were constructed from cell-line data that spanned all available cell types. Thus, we evaluated whether this heterogeneity may have confounded the analysis, leading to poor predictive performance. We repeated the training and testing steps on a cell-type specific basis. In cases with more than 5 cell lines for a given cell type, we trained models using only those cell lines and made predictions for TCGA samples associated with the same primary cell type. This too resulted in poor predictive performance (Table S24 and Figure S16), but the relatively small training sets were likely a limiting factor.

## Discussion

In an ideal setting, patient data would be used to train predictive models for clinical drug responses directly, as these data may accurately reflect tumor behavior in actual patients. Environmental factors, the tumor microenvironment, co-existing conditions, and a variety of other factors can affect a tumor’s behavior in ways that may not be accounted for in preclinical studies. However, acquiring drug-response data directly from human patients requires conducting many experimental tests on a given patient, which could be unethical, harmful, and subject to many confounding factors. In addition, patients are typically assigned standard-of-care protocols based on their specific cancer type. As a result, experimental drug-response data for large patient cohorts are scarcely available. An alternative approach is to use preclinical samples to identify molecular signatures of drug response and later use those signatures to predict clinical drug responses in patients.

Cell lines serve as preclinical models for drug development. Being able to accurately predict drug responses for a given cell line based on molecular features may help in optimizing drug-development pipelines and explain mechanisms behind treatment responses. We focused on DNA methylation profiles as one type of molecular feature that is known to drive tumorigenesis and modulate treatment responses (Esteller, 2002). When using classification or regression algorithms to predict discrete or continuous responses, respectively, we consistently observed excellent predictive performance when the training and test sets both consisted of cell-line data. Although conventional wisdom advises against discretizing a continuous response variable, where possible, due to loss of information, we wished to evaluate the potential to make effective predictions in this scenario, in part because clinical treatment responses are generally represented as discrete values. Using a subsampling approach, we observed that classification performance generally improved as more extreme examples were used for training and testing, whereas the opposite was true for the regression analyses. This suggests that during regression, the algorithms benefitted from seeing examples across a diverse range of IC_50_ values for a given drug, whereas the classification algorithms were confounded by seeing examples with relatively similar drug responses, even though sample sizes were smaller. This result has potential financial implications: if researchers can identify cell lines that are extreme responders for a particular drug, they may only need to generate costly molecular profiles for those cell lines. Future research may elucidate whether this finding generalizes to other types of molecular data and other drugs.

Previous efforts to associate DNA methylation levels with drug responses include work from Shen et al. (2007) who quantified methylation for 32 CpG islands in the NCI-60 cell lines, creating a sensitivity database for ~30k drugs and identifying biomarkers that predict drug sensitivity. Instead, our work uses microarray data to quantify methylation levels for thousands of genes across 987 cell lines but for fewer drugs. Rather than searching for individual genes that predict drug sensitivity, we constructed predictive models that represent patterns spanning as many as thousands of genes. Such an approach may better represent complex interactions among genes and thus yield improved predictive power, but a tradeoff is lack of model interpretability. We sought to shed some insight into the biological mechanisms that influence drug responses via feature selection, but methods for deriving such insights from genome-wide data are still in their infancy. Recent work using mathematical optimization models shows promise as a way to integrate molecular data from cell lines with drug-sensitivity information to infer resistance mechanisms (Fleck et al., 2016; Fleck et al., 2019).

A variety of computational methods have been proposed to predict drug responses for cell lines based on molecular data. Classical algorithms like decision trees and support vector machines have been used to predict the clinical efficiency of anti-cancer drugs and classify drug responses (Stetson et al., 2015; Borisov et al., 2018; Oskooei et al., 2018; Webber et al., 2018; Parca et al., 2019; Su et al., 2019). Neural networks (Menden et al., 2013) and deep neural networks (Chiu et al., 2019) have been used to predict drug response based on genomic profiles from cell lines. Other techniques have included elastic net regression (Basu et al., 2013; Webber et al., 2018; Parca et al., 2019), linear ridge regression (Geeleher et al., 2017), and LASSO regression (Huang et al., 2020). Alternative approaches based on computational linear algebra or network structures have also been applied to infer drug response in cell lines; these include matrix factorization (Guan et al., 2019), matrix completion (Nguyen and Le, 2018), and link prediction (Stanfield et al., 2017) methods. Finally, a community-based competition assessed the ability to predict therapeutic responses in cell lines using 44 regression-based algorithms (Costello et al., 2014). In our study we used diverse algorithms, but our primary focus was data subsampling and evaluating the potential to make accurate predictions of drug response in cell lines using relatively extreme responders.

We attempted to predict clinical responses for patients from TCGA, but the accuracy of these predictions was poor. Integrating datasets can introduce batch effects (Leek et al., 2010) and other systematic biases; we attempted to mitigate these biases using data that had been preprocessed identically for GDSC and TCGA. However, subtle differences in the way biological samples are handled and processed in the lab can make generalization difficult to achieve. Furthermore, inherent differences between cell lines and tumors may confound such predictions. Cell lines are grown in a controlled environment, and the cells are relatively homogeneous, whereas tumor samples are a heterogeneous milieu of cells. In addition, TCGA tumor responses were based on clinical observations instead of IC50 values, so there was no direct mapping between these types of measurements. Furthermore, our approach for quantifying predictive performance was different for the GDSC cross-validation analysis compared to the TCGA training/testing analysis. In the former, the class variable represented two possible outcomes (response and non-response). In the latter, the class variable was ordinal. Yet another challenge was that we used cell lines from all available cell types in GDSC. Better accuracy might be attained when training and testing on a single cell type; however, larger sample sizes would be necessary.

## Conclusion

We applied machine-learning algorithms to predict cytotoxic responses for eight anti-cancer drugs using genome-wide, DNA methylation profiles from 987 cell lines from the Genomics of Drug Sensitivity in Cancer (GDSC) database. We then compared the performance of classification and regression algorithms and evaluated the effect of sample size on model performance by artificially subsampling the data to varying degrees. We also used feature selection to identify informative genes with predictive power for drug response. For additional validation, we evaluated our ability to train a model based on drug responses in the GDSC cell lines and then accurately predict patient drug responses using data from The Cancer Genome Atlas (TCGA). Because patient-response values are categorical in nature, we only performed classification for these data.

Our study has limitations that could be addressed in future research. For one, we focus on DNA methylation profiles, but other types of molecular features contribute to tumorigenesis and modulate treatment responses. A number of cell-line studies have used gene-expression profiles to predict drug responses, and future studies could evaluate the potential benefits of incorporating more than one type of molecular feature into models for response prediction. Moreover, our analyses provide insights into the effect of subsampling on classification performance that have potential financial implications, but future research may elucidate whether these findings generalize to other types of molecular data and drugs. Finally, in our study, we discretize IC_50_ values in two classes (low IC_50_ values vs. high IC_50_ values), but TCGA data uses four classes to distinguish drug response (clinical progressive disease; stable disease; partial response; complete response). Additionally, treatment response data from patients is typically imbalanced, meaning that not all response classes contain similar numbers of patients. Hence, additional work could analyze the effect of class imbalance on model performance.

## Declarations

### Abbreviations

ACC: Accuracy
AUC: Area under the receiver operating characteristic curve
GDSC: Genomics of Drug Sensitivity in Cancer
KNN: K-nearest neighbors
MAE: Mean absolute error
MCC: Matthews correlation coefficient
MMCE: Mean misclassification error
MSE: Mean squared error
NB: NaÏve Bayes
RF: Random forests
RMSE: Root-mean-square deviation
SVM: Support vector machines
TCGA: The Cancer Genome Atlas
XGBoost: Extreme Gradient Boosting

## Acknowledgments

The authors express gratitude to patients who donated specimens and data to the GDSC and TCGA databases and those who curated the data and made it publicly available. We used computing resources from the Fulton Supercomputing Laboratory at Brigham Young University to perform these analyses.

## Authors’ contributions

All authors contributed to the study design. SPM performed all analyses and created all figures and tables. SPM, JLF, and SRP prepared the manuscript. FAB and PMM provided critical feedback on the manuscript.

## Funding

This work was supported in part by the National Council for Scientific and Technological Development (CNPq) under grant 428907/2018-0 (JLF); the Coordination for the Improvement of Higher Education Personnel (CAPES) - Finance Code 001; the Pontifical Catholic University of Rio de Janeiro.

## Availability of data and materials

Scripts for downloading and processing the data can be found at https://osf.io/r7pdb/.

## Ethics approval and consent to participate

Not applicable.

## Consent for publication

Not applicable.

## Competing interests

The authors declare that they have no competing interests.

## Supporting Information

**(Supplementary Figure) Figure S1: Example of subsampling process.** When performing classification, we discretized the drug-response (IC_50_) values. To evaluate alternative thresholds for discretization, we performed a subsampling analysis. In Scenario 1 illustrated above, we considered the cell lines with the lowest and highest 5% of IC_50_ values. In Scenario 2, we considered the cell lines with the lowest and highest 10% of IC_50_ values. Each scenario used 10% more data than the previous scenario (5% on each side). This pattern continues until all data were considered in the analysis.

**(Supplementary Figure) Figure S2: Graphs for Cisplatin classification analysis.** The graphs compare different scenarios ranked in order of best result. GDSC cell-line data were used to generate ten subsampling scenarios, which we then tested via nested cross validation. Scenarios that are further away from the center represent higher metric values than scenarios closer to it. The evaluated metrics for each algorithm are accuracy (ACC), specificity, recall, Matthews correlation coefficient (MCC), F1 score (F1) and area under the receiver operating characteristic curve (AUC).

**(Supplementary Figure) Figure S3: Graphs for Docetaxel classification analysis.** The graphs compare different scenarios ranked in order of best result. GDSC cell-line data were used to generate ten subsampling scenarios, which we then tested via nested cross validation. Scenarios that are further away from the center represent higher metric values than scenarios closer to it. The evaluated metrics for each algorithm are accuracy (ACC), specificity, recall, Matthews correlation coefficient (MCC), F1 score (F1) and area under the receiver operating characteristic curve (AUC).

**(Supplementary Figure) Figure S4: Graphs for Doxorubicin classification analysis.** The graphs compare different scenarios ranked in order of best result. GDSC cell-line data were used to generate ten subsampling scenarios, which we then tested via nested cross validation. Scenarios that are further away from the center represent higher metric values than scenarios closer to it. The evaluated metrics for each algorithm are accuracy (ACC), specificity, recall, Matthews correlation coefficient (MCC), F1 score (F1) and area under the receiver operating characteristic curve (AUC).

**(Supplementary Figure) Figure S5: Graphs for Etoposide classification analysis.** The graphs compare different scenarios ranked in order of best result. GDSC cell-line data were used to generate ten subsampling scenarios, which we then tested via nested cross validation. Scenarios that are further away from the center represent higher metric values than scenarios closer to it. The evaluated metrics for each algorithm are accuracy (ACC), specificity, recall, Matthews correlation coefficient (MCC), F1 score (F1) and area under the receiver operating characteristic curve (AUC).

**(Supplementary Figure) Figure S6: Graphs for Gemcitabine classification analysis.** The graphs compare different scenarios ranked in order of best result. GDSC cell-line data were used to generate ten subsampling scenarios, which we then tested via nested cross validation. Scenarios that are further away from the center represent higher metric values than scenarios closer to it. The evaluated metrics for each algorithm are accuracy (ACC), specificity, recall, Matthews correlation coefficient (MCC), F1 score (F1) and area under the receiver operating characteristic curve (AUC).

**(Supplementary Figure) Figure S7: Graphs for Paclitaxel classification analysis.** The graphs compare different scenarios ranked in order of best result. GDSC cell-line data were used to generate ten subsampling scenarios, which we then tested via nested cross validation. Scenarios that are further away from the center represent higher metric values than scenarios closer to it. The evaluated metrics for each algorithm are accuracy (ACC), specificity, recall, Matthews correlation coefficient (MCC), F1 score (F1) and area under the receiver operating characteristic curve (AUC).

**(Supplementary Figure) Figure S8: Graphs for Temozolomide classification analysis.** The graphs compare different scenarios ranked in order of best result. GDSC cell-line data were used to generate ten subsampling scenarios, which we then tested via nested cross validation. Scenarios that are further away from the center represent higher metric values than scenarios closer to it. The evaluated metrics for each algorithm are accuracy (ACC), specificity, recall, Matthews correlation coefficient (MCC), F1 score (F1) and area under the receiver operating characteristic curve (AUC).

**(Supplementary Figure) Figure S9: Graphs for Gefitinib regression analysis.** We used DNA methylation data from cell lines to predict continuous IC_50_ response values using four regression algorithms. We evaluated the algorithms’ performance via nested cross validation for ten subsampling scenarios. The graphs illustrate the performance for these scenarios, ranked in order of relative performance for three metrics: MSE (Mean Squared Error), RMSE (Root Mean Square Error), and MAE (Mean Absolute Error). Scenarios further away from the center represent relatively low metric values (and thus better performance). Scenarios that used all cell lines performed best for all algorithms.

**(Supplementary Figure) Figure S10: Graphs for Docetaxel regression analysis.** We used DNA methylation data from cell lines to predict continuous IC_50_ response values using four regression algorithms. We evaluated the algorithms’ performance via nested cross validation for ten subsampling scenarios. The graphs illustrate the performance for these scenarios, ranked in order of relative performance for three metrics: MSE (Mean Squared Error), RMSE (Root Mean Square Error), and MAE (Mean Absolute Error). Scenarios further away from the center represent relatively low metric values (and thus better performance). Scenarios that used all cell lines performed best for all algorithms.

**(Supplementary Figure) Figure S11: Graphs for Doxorubicin regression analysis.** We used DNA methylation data from cell lines to predict continuous IC_50_ response values using four regression algorithms. We evaluated the algorithms’ performance via nested cross validation for ten subsampling scenarios. The graphs illustrate the performance for these scenarios, ranked in order of relative performance for three metrics: MSE (Mean Squared Error), RMSE (Root Mean Square Error), and MAE (Mean Absolute Error). Scenarios further away from the center represent relatively low metric values (and thus better performance). Scenarios that used all cell lines performed best for all algorithms.

**(Supplementary Figure) Figure S12: Graphs for Etoposide regression analysis.** We used DNA methylation data from cell lines to predict continuous IC_50_ response values using four regression algorithms. We evaluated the algorithms’ performance via nested cross validation for ten subsampling scenarios. The graphs illustrate the performance for these scenarios, ranked in order of relative performance for three metrics: MSE (Mean Squared Error), RMSE (Root Mean Square Error), and MAE (Mean Absolute Error). Scenarios further away from the center represent relatively low metric values (and thus better performance). Scenarios that used all cell lines performed best for all algorithms.

**(Supplementary Figure) Figure S13: Graphs for Gemcitabine regression analysis.** We used DNA methylation data from cell lines to predict continuous IC_50_ response values using four regression algorithms. We evaluated the algorithms’ performance via nested cross validation for ten subsampling scenarios. The graphs illustrate the performance for these scenarios, ranked in order of relative performance for three metrics: MSE (Mean Squared Error), RMSE (Root Mean Square Error), and MAE (Mean Absolute Error). Scenarios further away from the center represent relatively low metric values (and thus better performance). Scenarios that used all cell lines performed best for all algorithms.

**(Supplementary Figure) Figure S14: Graphs for Paclitaxel regression analysis.** We used DNA methylation data from cell lines to predict continuous IC_50_ response values using four regression algorithms. We evaluated the algorithms’ performance via nested cross validation for ten subsampling scenarios. The graphs illustrate the performance for these scenarios, ranked in order of relative performance for three metrics: MSE (Mean Squared Error), RMSE (Root Mean Square Error), and MAE (Mean Absolute Error). Scenarios further away from the center represent relatively low metric values (and thus better performance). Scenarios that used all cell lines performed best for all algorithms.

**(Supplementary Figure) Figure S15: Graphs for Temozolomide regression analysis.** We used DNA methylation data from cell lines to predict continuous IC_50_ response values using four regression algorithms. We evaluated the algorithms’ performance via nested cross validation for ten subsampling scenarios. The graphs illustrate the performance for these scenarios, ranked in order of relative performance for three metrics: MSE (Mean Squared Error), RMSE (Root Mean Square Error), and MAE (Mean Absolute Error). Scenarios further away from the center represent relatively low metric values (and thus better performance). Scenarios that used all cell lines performed best for all algorithms.

**(Supplementary Figure) Predicting patient drug response from methylation profiles for the LGG cell type and Temozolomide drug (n=79).** For each TCGA test sample, our model (+-10%c SVM) predicted probabilities to drug response based only on LGG cell-type. However, the classification algorithm was not able to identify active patterns to translate methylation profile tumor drug response prediction.

**(Supplementary Table) Table S1: Classification results for all combinations of subsampling scenarios and algorithms for Cisplatin**.

**(Supplementary Table) Table S2: Classification results for all combinations of subsampling scenarios and algorithms for Docetaxel**.

**(Supplementary Table) Table S3: Classification results for all combinations of subsampling scenarios and algorithms for Doxorubicin**.

**(Supplementary Table) Table S4: Classification results for all combinations of subsampling scenarios and algorithms for Etoposide**.

**(Supplementary Table) Table S5: Classification results for all combinations of subsampling scenarios and algorithms for Gemcitabine**.

**(Supplementary Table) Table S6: Classification results for all combinations of subsampling scenarios and algorithms for Paclitaxel**.

**(Supplementary Table) Table S7: Classification results for all combinations of subsampling scenarios and algorithms for Temozolomide**.

**(Supplementary Table) Table S8: Regression results for all combinations of subsampling scenarios and algorithms for Cisplatin**.

**(Supplementary Table) Table S9: Regression results for all combinations of subsampling scenarios and algorithms for Docetaxel**.

**(Supplementary Table) Table S10: Regression results for all combinations of subsampling scenarios and algorithms for Doxorubicin**.

**(Supplementary Table) Table S11: Regression results for all combinations of subsampling scenarios and algorithms for Etoposide**.

**(Supplementary Table) Table S12: Regression results for all combinations of subsampling scenarios and algorithms for Gemcitabine**.

**(Supplementary Table) Table S13: Regression results for all combinations of subsampling scenarios and algorithms for Paclitaxel**.

**(Supplementary Table) Table S14: Regression results for all combinations of subsampling scenarios and algorithms for Temozolomide**.

**(Supplementary Table) Table S15: Informative genes for predicting cell-line responses for Cisplatin.** We used the feature selection to identify informative genes for Cisplatin drug-response prediction. Genomic coordinates are based on build 37 of the human genome. We used information gain to rank the genes; a higher score indicates a more informative gene.

**(Supplementary Table) Table S16: Informative genes for predicting cell-line responses for Docetaxel.** We used the feature selection to identify informative genes for Docetaxel drug-response prediction. Genomic coordinates are based on build 37 of the human genome. We used information gain to rank the genes; a higher score indicates a more informative gene.

**(Supplementary Table) Table S17: Informative genes for predicting cell-line responses for Doxorubicin.** We used the feature selection to identify informative genes for Doxorubicin drug-response prediction. Genomic coordinates are based on build 37 of the human genome. We used information gain to rank the genes; a higher score indicates a more informative gene.

**(Supplementary Table) Table S18: Informative genes for predicting cell-line responses for Etoposide.** We used the feature selection to identify informative genes for Etoposide drug-response prediction. Genomic coordinates are based on build 37 of the human genome. We used information gain to rank the genes; a higher score indicates a more informative gene.

**(Supplementary Table) Table S19: Informative genes for predicting cell-line responses for Gemcitabine.** We used the feature selection to identify informative genes for Gemcitabine drug-response prediction. Genomic coordinates are based on build 37 of the human genome. We used information gain to rank the genes; a higher score indicates a more informative gene.

**(Supplementary Table) Table S20: Informative genes for predicting cell-line responses for Paclitaxel.** We used the feature selection to identify informative genes for Paclitaxel drug-response prediction. Genomic coordinates are based on build 37 of the human genome. We used information gain to rank the genes; a higher score indicates a more informative gene.

**(Supplementary Table) Table S21: Informative genes for predicting cell-line responses for Temozolomide.** We used the feature selection to identify informative genes for Temozolomide drug-response prediction. Genomic coordinates are based on build 37 of the human genome. We used information gain to rank the genes; a higher score indicates a more informative gene.

**(Supplementary Table) Table S22: Gene-set evaluation for the classification analysis**. We used a statistical overrepresentation test to identify protein classes associated with the top-20 ranked genes in the feature-selection analysis.

**(Supplementary Table) Table S23: Gene-set evaluation for the regression analysis.** We used a statistical overrepresentation test to identify protein classes associated with the top-20 ranked genes in the feature-selection analysis.

**(Supplementary Table) Table S24: Slope values for viable combinations of subsampling scenarios, algorithms, drug type and primary cell type.** Training and testing stages were evaluated on a cell-type specific basis if the number of existing cell lines was greater than 5 for a given cell type. Four models (+-5%c SVM, +-5%c RF, +-10%c SVM and +-10%c RF) were trained using only those cell lines and made predictions for the associated TCGA samples. Slopes close to zero show that the evaluated models were unable to identify patterns that successfully translated into patient responses. Tissue acronyms: BLCA=Urothelial Bladder Carcinoma, BRCA=Breast Invasive Carcinoma, CESC=Cervical Squamous Cell Carcinoma and Endocervical Adenocarcinoma, GBM=Glioblastoma Multiforme, HNSC= Head-Neck Squamous Cell Carcinoma, LGG=Low Grade Glioma, LUAD=Lung Adenocarcinoma, LUSC= Lung Squamous Cell Carcinoma, SKCM= Skin Cutaneous Melanoma, STAD= Stomach Adenocarcinoma, UCEC= Uterine Corpus Endometrial Carcinoma

